# Identification of the stress granule transcriptome via RNA-editing in single cells and *in vivo*

**DOI:** 10.1101/2021.06.21.449212

**Authors:** Wessel van Leeuwen, Michael VanInsberghe, Nico Battich, Fredrik Salmén, Alexander van Oudenaarden, Catherine Rabouille

## Abstract

Stress granules are phase separated assemblies formed around mRNAs whose identities remain elusive. The techniques available to identify the RNA content of stress granules rely on their physical purification, and are therefore not suitable for single cells and tissues displaying cell heterogeneity. Here, we adapted TRIBE (Target of RNA-binding proteins Identified by Editing) to detect stress granule RNAs by fusing a stress granule RNA-binding protein (FMR1) to the catalytic domain of an RNA-editing enzyme (ADAR). RNAs colocalized with this fusion are edited, producing mutations that are detectable by sequencing. We first optimized the expression of this fusion protein so that RNA editing preferentially occurs in stress granules. We then show that this purification-free method can reliably identify stress granule RNAs in bulk and single S2 cells, and in Drosophila tissues, such as 398 neuronal stress granule mRNAs encoding ATP binding, cell cycle and transcription factors. This new method opens the possibility to identify the RNA content of stress granules as well other RNA based assemblies in single cells derived from tissues.

## INTRODUCTION

Non-membrane bound compartments represent an important aspect of cell organization. They are formed by phase separation where solution of seemingly diffuse macromolecules segregate into two distinct phases (1,2). Interestingly, cellular stress induces the formation of many of these compartments (3), including stress granules (4,5).

Stress granules are one of these phase-separated assemblies that formed around mRNAs after cells have been subjected to cellular stress leading to inhibition of protein translation initiation and to polysome disassembly (6). This results in ribosome-free accumulation of untranslated mRNAs in the cytoplasm and their binding to RNA-binding proteins (RBPs) that coalesce into membraneless foci, the stress granules (7).

Over the years, the protein content of stress granules has been extensively studied (5,8–10). In particular, RBPs such as G3BP1/2 (11) and TIA1 (12) have been shown to be critical for the formation of stress granules, but many others proteins are also present in these granules (the so-called “clients”), such as Caprin (13) and FMR1 (14). In the meantime, the biophysical principles underlying stress granule phase separation has been elucidated in vitro using purified RBPs and their RNA binding domains (15–17)

Recently, however, RNAs have also been shown to be structural components of stress granules (18,19) both in vitro and in vivo. This is sustained by the demonstration that elements in RNA secondary structure are conducive to phase separation into liquid/solid phase separated droplets (20). This renders the identification of stress granule RNAs an important biological question.

The identification of stress granule RNAs has recently been performed in mammalian cultured cells after stress granule core isolation (21,22), cross-linking and immunoprecipitation (23), as well as proximity biotinylation of RNAs (24). These studies revealed that stress granules mainly harbor long mRNAs that are thought to be poorly translated by ribosomes. Interestingly, although the cell types used in the two of these methods is different (21,22), they exhibited a 67 % overlap in RNA species, indicating a significant degree of conservation. Furthermore, even though a clear enrichment in adenylate-uridylate AU-rich elements (ARE) has been established in ER-stress driven stress granules mRNAs (21), a true recruitment motif has not yet been identified. Recent efforts shows that AREs and Pumilio recognition elements increase mRNA partitioning into stress granules (25).

However valuable, these techniques are somewhat limited by the step of cell fractionation and stress granules purification. Indeed, these include the risk of losing materials (8,26), and they require a large amount of input material, thus preventing the analysis of limited tissues and single cells. This represents a critical step to compare the differential response to stress of a given cell in its environment. It varies from stress to stress (5), and there are now indications that the RNA content of stress granules might be heterogeneous within a cell population (22). In this regard, it is widely proposed that RNAs are recruited in stress granules to be protected from degradation by cytoplasmic nucleases during stress, and are readily available to re-enter translation upon stress relief, thus giving the cells a fitness advantage. The identification of RNA species present in stress granules in single cells is therefore relevant to address this hypothesis and predict whether a cell would thrive after a period of stress.

Here, we developed a method to identify RNAs that are recruited to stress granules, which does not require fractionation and isolation, and that works in single cells and in tissue in a cell type specific manner. This method is based on TRIBE (Target of RNA-binding proteins Identified by Editing) by the Rosbash group (27). There, a given RBP is fused to the catalytic domain of the DNA/RNA editing enzyme ADAR (ADARcd) that deaminates adenosine-to-inosine on RNA molecules, which is read out as A-to-G mutations by sequencing. This method has been used to successfully identify the RNA targets of several RPBs in *Drosophila* neurons (27). Furthermore, a hyperactivating mutation in ADARcd (HyperTRIBE) has been shown to increase the editing efficiency of ADARcd by 400% when compared to the original ADARcd (28).

The purification-free method we developed has allowed for the first time the identification of stress granule RNAs in single cells, and in specific tissues, here the neurons of the *Drosophila* larval brain.

## MATERIAL AND METHODS

### Experimental model

Wild type *Drosophila* S2 cells were cultured in Schneider’s medium (Sigma Aldrich) supplemented with 10 % insect tested fetal bovine serum (Sigma Aldrich) at 26 °C. This medium is referred to as Schneider’s. *Drosophila* S2 cells stably transfected with pMT-FMR1-ADARcd-V5 (see below) were grown in Schneider’s supplemented with 300 μg/mL hygromycin B (ThermoFisher).

Flies were reared on rich fly food (https://bdsc.indiana.edu/information/recipes/germanfood.html) and maintained at 25 °C with 70 % humidity with a 12 h light/dark cycle.

### Plasmid construction and generation of stable S2 clones and transgenic flies

The plasmid used to express FMR1-ADARcd in S2 cells was pMT-FMR1-ADARcd-V5. This plasmid was generated by amplifying hyperactive ADARcd (that contains the hyperactivating E488Q mutation (28) from a gblock (IDT) containing ADARcd using the primers 5’-CGTATCGTCTCGAGATAC-3’ (forward) and 5’-GTACGTTAGTTTAAACTCACG-3’ (reverse). This was ligated into the plasmid pMT-FMR1-V5 (29) using the restriction enzymes XhoI (NEB) and PmeI (NEB). To obtain stable clones, pMT-FMR1-ADARcd-V5 was co-transfected with pHygro in S2 cells using Effectene transfection reagent (Qiagen) with a 1:10 ratio of DNA to Effectene Reagent. Stable S2 cells were generated by culturing cells in 300 μg/mL Hygromycin B (ThermoFisher).

To generate transgenic flies expressing FMR1-ADARcd-V5, we generate the plasmid pUAS-FMR1-ADARcd-V5-T2A-EGFP by Infusion (TaKaRa) cloning of the three PCR fragments, FMR1-ADARcd-V5, 2x T2A, and EGFP into the plasmid pUAS. FMR1-ADARcd-V5 was amplified out of pMT-FMR1-ADARcd-V5 using the primers 5’-TGAATAGGGAATTGGGAATTCACAAAATGGAAGATCTCCTCGTGGAAGT-3’ (forward) and 5’-CCAGATCTCCCGTAGAATCGAGACCGAGGAGAG-3’ (reverse). 2x T2A was amplified out of pMT-2xT2A using primers 5’-CGATTCTACGGGAGATCTGGGCAGTGGAGAGG-3’ (forward) and 5’-TGCTCACCATCTCGAGCCCGGGTCCT-3’ (reverse). EGFP was amplified out pEGFP (13031, Addgene) using primers 5’-CGGGCTCGAGATGGTGAGCAAGGGCGAGGAG-3’ (forward) and 5’-TTCACAAAGATCCTCTAGAGGTACCTCACTTGTACAGCTCGTCCATGCCG-3’ (reverse). Transgenic flies were generated by BestGene using P-element based services and balanced with CyO or TM6. All primers are documented in *Suppl Table S4*.

### Arsenite treatment

Wild type *Drosophila* S2 cells or *Drosophila* S2 cells stably transfected with pMT-FMR1-ADARcd-V5 were plated to a concentration of 1.5 × 10^6^ cells per well in a 12-wells plate containing a coverslip and allowed to adhere for 1 h at 26 °C in Schneider’s. The expression of FMR1-ADARcd-V5 was induced by addition of 1 mM CuSO_4_ for 4 h at 26 °C. This was followed by either a further incubation in Schneider’s (basal condition) or incubation with 500 μM NaAsO_2_ (arsenite) for 4 h at 26 °C.

### Induction geneswitch, isolation and dissociation *Drosophila* brain, and arsenite treatment

To induce the expression of FMR1-ADARcd-V5-T2A-EGFP specifically in neurons, we crossed homozygous UAS-FMR1-ADARcd-V5-T2A-EGFP male flies (on second chromosome) with homozygous elav-geneswitchGal4 (43642, BDSC) female flies (on third chromosome). Eggs were laid in large tubes containing rich fly food. After two days, flies were removed and the tubes were further incubated for 3 days at 25 °C. 30 wandering third instar UAS-FMR1-ADARcd-V5-T2A-EGFP; *elav*-GeneSwitch-Gal4 larvae were collected in a 2 mL Eppendorf tube, washed 2 times with MiliQ, washed once with 80 % ethanol, and incubated in 3 mg/mL RU486 (in 80 % ethanol, M8046, Sigma Aldrich) for 10 min to activate the geneswitch. Larvae were transferred back to a fresh apple juice plate and allowed to recover for 14 h in the dark at RT (20 °C). At least 20 third instar UAS-FMR1-ADARcd-V5-T2A-EGFP; *elav*-GeneSwitch-Gal4 larvae were collected, washed once with PBS, and dissected. Their brains were isolated in ice cold Schneider’s media in the lid of 6 cm dish and transferred to an Eppendorf tube containing ice cold Schneider’s. To dissociate the brains, the isolated brains were washed 2 times with Rinaldini solution (30), and incubated with 1 mg/mL collagenase I and papain (Sigma Aldrich) in Rinaldini solution for 1 h at 30 °C in a thermomixer. Schneider’s was then added and brains were washed 2 times with Schneider’s on ice. Brains were disrupted manually by pipetting up and down 75 times with a 200 μL pipet tip. The tissue pieces were forced through a cell-strainer FACS tube (BD Falcon) and either plated on poly-L-lysin (P1274, Sigma Aldrich) coated coverslips for immunofluorescence (IF) and single molecule FISH (smFISH), or in a well of a 12 well plates for RNA isolation (see below). Upon adhesion for 1 h at RT in the dark (20 °C), the neurons were either incubated in Schneider’s or stressed with 500 μM NaAsO_2_ for 4 h at 26 °C.

### Antibodies

For immunofluorescence, we used the primary antibodies mouse-anti-dFMR1 (1:20) (5A11, DSHB), Mouse-anti-caprin (1:100) (31), Mouse-anti-V5 (1:500) (R960-25, Thermofisher) and Rabbit-anti-V5 (1:500) (V8137, Sigma Aldrich), Rat-anti-elav (1:20) (7E8A10, DSHB), Mouse-anti-Repo (1:30) (8D12, DSHB), Goat-anti-mouse Alexa 488 (A11001, Invitrogen), Donkey-anti-Rabbit Alexa 568 (A10042, Invitrogen), Donkey-anti-mouse Alexa 568 (A10037, Invitrogen), Donkey-anti-mouse Alexa 647 (A31571, Invitrogen), Goat-anti-Rat Alexa 633 (A21094, Invitrogen) and Donkey-anti-Rabbit Alexa 647 (A31573, Invitrogen) as secondary antibodies.

For Western blot, we used a mouse-anti-V5 (1:500) (R960-25, Thermofisher) and mouse-anti-tubulin (1:10000) (T5168, Sigma Aldrich) as primary antibody. Sheep-anti-mouse coupled to HRP (1:2000) (GENA931, GE Healthcare) was used as secondary antibody.

### Immunofluorescence (IF)

After treatments, S2 cells and dissociated brain cells were fixed with 4 % paraformaldehyde (Sigma Aldrich) in PBS (pH 7.4) for 20 min. Cells were then washed 3 times with PBS and subsequently quenched by incubation in 50 mM NH_4_Cl in PBS for 5 min. Followed by permeabilization with 0.11 % Triton-X for 5 min. Hereafter, cells were washed 3 times in PBS and blocked in PBS supplemented with 0.5 % fish skin gelatin (Sigma Aldrich) for 20 min. Cells were then incubated with the primary antibody (in blocking buffer) for 25 min, washed 3 times with blocking buffer and incubated with the secondary antibody (in blocking buffer) coupled to a fluorescent dye for 20 min. Cells on the coverslip were washed 2 times in milliQ and dried for 3 min on a tissue with cells facing up. Finally, each coverslip with cells was mounted with Prolong antifade media (+DAPI)(ThermoFisher) on a microscope slide.

### Western blot of FMR1-ADARcd-V5

A total of 3 × 10^6^ S2 cells expressing pMT-FMR1-ADARcd-V5 for 4 h were plated in a 6-wells plate per condition. They were harvested on ice and lysed in the lysis buffer containing 50 mM Tris-HCl pH 7.5, 150 mM NaCl, 1 % Triton X-100, 50 mM NaF, 1 mM Na_3_VO_4_, 25 mM Na_2_-β-glycerophosphate supplemented with a protease inhibitor tablet (Roche). The lysates were cleared by centrifugation at 14,000 rpm for 20 min at 4 °C. Supernatants were collected and the protein concentration was determined using the Pierce BCA protein assay kit (ThermoFisher). 20 µg of protein was mixed with 5× SDS loading dye, boiled for 5 min and fractionated on an 8 % SDS-PAGE gel. Fractionated proteins were transferred to a polyvinylidene difluoride (PVDF) membrane. After blocking in TBS+0.05 %Tween-20 supplemented with 5 % BSA (Sigma Aldrich) (Blocking buffer), primary antibodies (anti-V5 to detect FMR1-ADARcd-V5 and alpha tubulin) were added for an overnight incubation at 4 °C. The membrane was subsequently washed 3 times in TBS+0.05 % Tween-20 over 45 min and incubated with secondary antibodies for 1 h at RT (20 °C) after washing 3 times in TBS+0.05 %Tween-20 and developed by enhanced chemiluminescence (Bio-Rad) with Image Quant™ LAS 4000.

### Single Molecule FISH

Plated Wildtype S2 cells were first treated with arsenite (see above), fixed and labeled for endogenous FMR1 as described above in the IF section. After incubation with the secondary antibody, cells were washed 3 times with PBS and cells were post-fixed in 4 % paraformaldehyde in PBS (pH 7.4) for 10 min. Following a washing 3 times in PBS, cells were further incubated for 5 min in 10 % formamide (Thermofisher) in DEPC-treated water. They were then incubated overnight on a droplet containing one fluorescent smFISH probe (125 nM in 1 % dextransulfate (D8906, Sigma Aldrich), 10 % formamide (Thermofisher) in DEPC-treated water at 37 °C) in a moistened chamber to avoid drying. Cells were washed 2 times for 30 min with 10 % formamide (Thermofisher) in DEPC-treated water and mounted with Prolong antifade media (+DAPI) (ThermoFisher) on a microscope slide. All smFISH probes (*Suppl Table S4*) were labeled with Atto565 following (32). The DNA oligos were purchased from IDT. The TMR-oligo(dT)30x was also purchased from IDT.

### Microscopy and image analysis

Immunofluorescence images were acquired on a TSC SP8 confocal microscope (Leica GmbH) with a 63× lens. For smFISH, a widefield Leica MM-AF microscope was used with a 100× lens. smFISH images were deconvoluted using the Regularized Inverse Filter algorithm with the DeconvolutionLab2 plugin (33) in ImageJ. The minimum and maximum display values were adjusted using the Imaris Image Analysis Software (Bitplane) for each channel. The smFISH spots (each corresponding to a single RNA molecule, (34)) were counted using the spot count tool, and the percentage of spots in stress granules was estimated by their colocalization with FMR1.

### Treatment and isolation of RNA from S2 cells and dissociated brain cells

*Drosophila* S2 cells stably transfected with pMT-FMR1-ADARcd-V5 were plated to a concentration of 3 × 10^6^ cells per well in a 6-wells plate 16 h before the experiment. The expression of FMR1-ADARcd-V5 was induced for 4 h followed by arsenite treatment as described above. The dissociated brain cells came from 20 third instar larval brains and were treated as above. The media was removed and Trizol (Zymo Research) was added to lyse cells. RNA was extracted from the Trizol using the Direct-zol RNA microprep kit (Zymo Research) following manufacturer’s instructions. The RNA concentration was determined using the Qubit RNA High Sensitivity assay kit (Thermofisher). The quality of the RNA was analyzed by running the samples on an RNA 6000 Pico chip (Agilent Technologies) on the Bioanalyzer (Agilent Technologies).

### FACS – single-cell sorting

S2 cells were harvested without the requirement of dissociation (S2 cells grow as single cells). Single-S2-cell suspensions were resuspended in PBS0 with 1 µg/mL DAPI, and passed through a 20-µm mesh. Single cells were index sorted using a BD FACS Influx into 384-well hardshell plates (BioRad) that were pre-filled with 5 µL of light mineral oil (Sigma Aldrich) and 50 nL of 0.25 µM CelSeq2 primer (CS2001 to CS2384, *Suppl Table S4*) (35). Doublets, debris, and dead cells were excluded by gating forward and side scatter in combination with the DAPI channel.

### Library preparation and Sequencing

Library preparation was performed similar to sort-seq (35). Quality of the final libraries was assessed using the Agilent High Sensitivity DNA bioanalyzer (Agilent) and quantified with the Qubit (Invitrogen) before sequencing. Libraries were sequenced using v2.5 chemistry on a NextSeq 500 (Illumina).

### Data Analysis – Reference genomes and annotations

The Drosophila reference genome (BDGP6), annotations, and known variants were obtained from Ensembl release 95. Ribosomal RNA sequences used for read depletion were downloaded from FlyBase. The reference genomes were prepared for alignment by masking all tRNA genes and pseudogenes and including unique mature tRNAs genes as artificial chromosomes. tRNA genes and pseudogenes were identified using tRNAscan-SE (version 2.0.7) (36) using the eukaryotic model (-HQ) and the vertebrate mitochondrial model (-M vert -Q).

### Data Analysis – Read processing

Adapter and homopolymer sequences were first trimmed from read 2 using cutadapt (version 3.2) (37). Next, trimmed reads were depleted of rRNA sequences by aligning to a ribosomal reference using bwa aln (version 0.7.17-r1188) and bwa mem (38), and discarding any read that aligned using either method. Depleted reads were then two-pass aligned to the reference genome using STARSolo (version 2.7.7a) (39) with a 125-nt overhang. Aligned reads were deduplicated with UMI-tools (version 1.1.1) (40) using --spliced-is-unique --per-gene --per-cell.

### Data Analysis – Variant identification

Variant identification followed the GATK RNA-seq short variant discovery best-practices workflow using GATK (version 4.1.9.0) (41). First, the alignments were merged, and then reads with N in the alignment cigar were split into multiple alignments (SplitNCigarReads). Next, there were two rounds of base quality score recalibration (BaseRecalibrator, ApplyBQSR, and AnalyzeCovariates) and variant calling (HaplotypeCaller). The first round of recalibration masked all known reference variants, and the second additionally masked all novel variants. The exonic variants from the second round of variant calling were used for further downstream analyses.

### Data Analysis – Statistical tests

To identify the RNAs differentially more edited upon stress conditions, we set that the editing frequency of RNAs from the “arsenite” triplicates should be significantly higher (p<0.01 using the Empirical Bayes test (ebbr package in R)) than the editing frequency of same RNAs from the “basal triplicate”. The correlation (R2) between samples was calculated using the Python package pandas.dataframe.corr(). A student’s T-test (scipy.ttest) was used to determine the significance between mRNA length.

## RESULTS

### Predictions and experimental setup

To identify the RNA content in stress granules, we adapted the TRIBE method (27,28) in such a way that most of the RNA editing takes place in stress granules. The prediction is that the more an RNA is specifically edited in stress conditions (here arsenite treatment), the more likely it is recruited in stress granules. As in TRIBE, we used FMR1-ADARcd-V5 because endogenous and GFP tagged FMR1 are actively recruited to stress granules upon arsenite stress, making it likely that FMR1-ADARcd-V5 would behave similarly. Once recruited and concentrated it would edit the RNAs that are present in these assemblies **(Figure 1A)**. These edited RNAs will include FMR1 clients, but we hypothesize that they will also include RNAs that are clients of other stress granule RBPs and localised in close proximity, thus, edited in trans.

**Figure 1:**
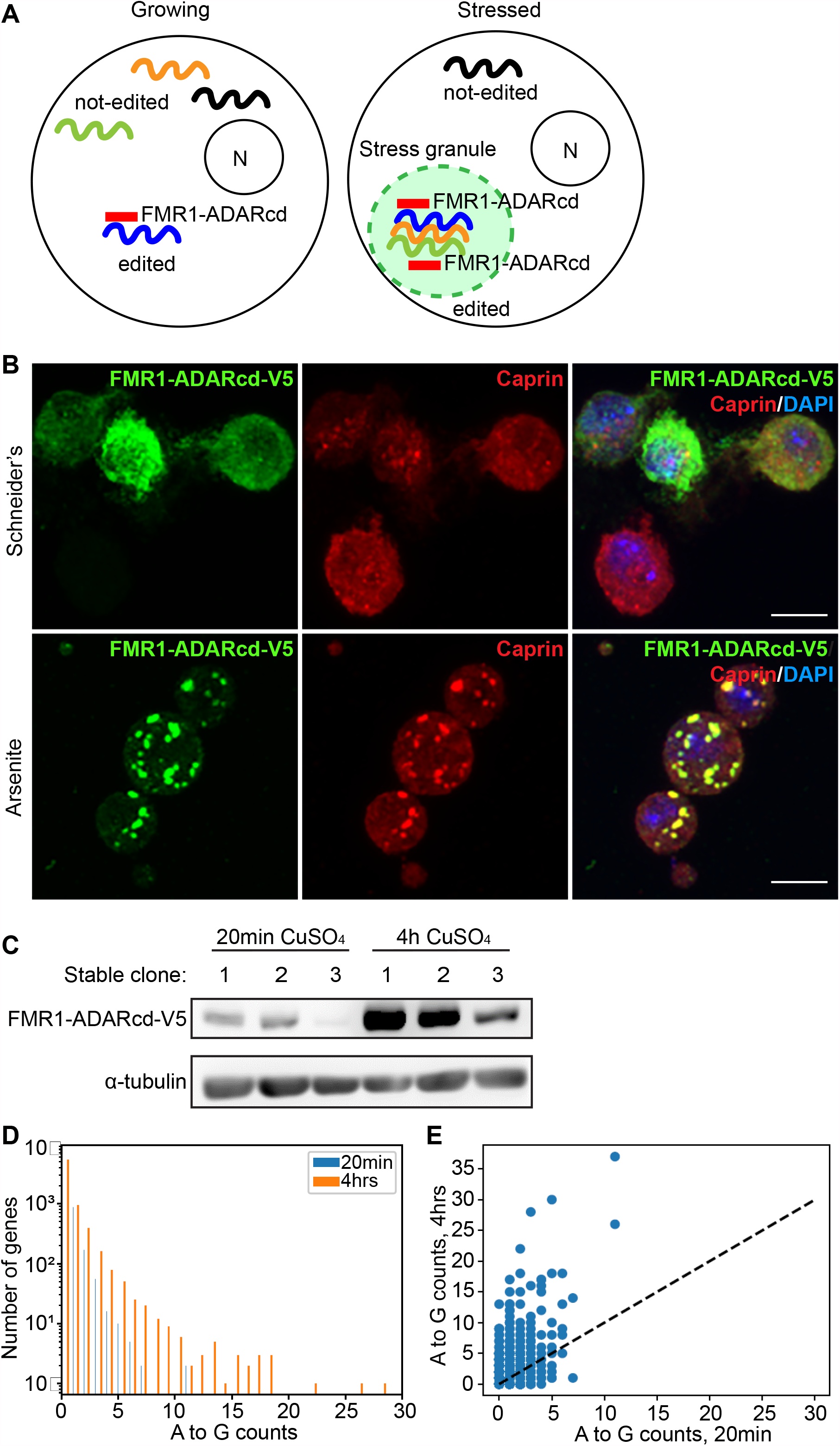
Adaptation of TRIBE using FMR1-ADARcd-V5 to detect stress granules RNAs. **A:** Schematic overview of our TRIBE adaptation strategy using FMR1-ADARcd-V5 to detect stress granules RNAs. In growing cells, FMR1-ADARcd-V5 is seemingly diffuse and binds to its own client RNAs that can be edited. Other non-client RNAs are not edited. Upon arsenite stress, many RNA-binding proteins are recruited in stress granules form, including FMR1-ADARcd-V5. This allows ADARcd to edit all the stress granules RNAs, potentially the FMR1 clients and other stress granules RNAs that in close proximity. The prediction is that the close proximity of ADARcd and RNAs increases their editing. It follows RNAs that are preferentially edited upon arsenite stress are likely to be located in stress granules. **B:** Immunofluorescence staining of FMR1-ADARcd-V5 in S2 cells upon incubation in Schneider’s and in Schneider’s + 0.5mM arsenite for 4 h. Upon arsenite, FMR1-ADARcd-V5 colocalizes with the known stress granule marker Caprin. **C:** Western blot of S2 cell extract after inducing the expression of FMR1-ADARcd-V5 with coppersulfate for 20 min or 4 h in different clones. Note that inducing the expression for 4 h leads to the visible expression of the chimeric protein in clone 1. **D:** Graph displaying the frequency of A>G editing events in S2 cells in which the expression of FMR1-ADARcd-V5 was induced for 20 min or 4 h. Note that the number of genes that are edited and the A>G editing frequency is higher after 4 h of induction. **E:** Graph displaying the total A>G editing events in S2 cells per gene after 20 min or 4 h of induction of FMR1-ADARcd-V5. The number of edits per gene is higher after 4 h. Scale bar: 10 μm (B)

To achieve this, we first checked that FMR1-ADARcd-V5 is indeed efficiently recruited in arsenite induced stress granules. We generated S2 cell clones stably expressing the FMR1-ADARcd-V5 fusion protein (HyperTRIBE) and found that as expected, it is efficiently recruited in foci upon arsenite treatment where it colocalized with the RNA-binding protein Caprin, a known marker of stress granules, in 90% of the cells (**Figure 1B**).

Second, for our prediction to be true, i.e. to detect a significant differential RNA editing in stress conditions, the RNA editing in basal conditions (non-stressed cells during which FMR1-ADARcd-V5 can edit FMR1 clients independently of their further recruitment in stress granules) should be kept as low as possible. The notion was to find a tradeoff in producing enough of chimeric protein to lead to RNA editing in stress granules without generating a high background in non-stressed conditions. To do so, we expressed FMR1-ADARcd-V5 (that is under control of a copper sulfate inducible metallothionein promoter) for 20 min and 4 h (instead of 24 h as in (27), before applying the stress. Both induction times led to detectable FMR1-ADARcd-V5 protein (**Figure 1C)**. We chose clone 1 for the rest of the experiments.

We then verified that the induction of FMR1-ADARcd-V5 is able to produce detectable editing events in non-stressed conditions. Calling variants on these samples (42) identified 1,727 RNAs with modified bases. As expected, most of the filtered de-novo variants detected were A-to-G events (**Figure D**), while other possible nucleotide changes were hardly detected (*Suppl figure S1A-C*). Interestingly, comparing the levels of RNA editing of the 20 min to the 4 h induction time showed clear evidence of an increased RNA editing (**Figure 1D, E**). However, we chose 4 h induction time to get sufficient amount of protein that leads to editing.

### RNA editing through ADAR predicts the stress granule transcriptome in S2 cells

To identify RNAs present in stress granules, we used the setup conditions described above (4 h induction) followed by cell incubation with arsenite for 4 h. Control cells were maintained in Schneider’s for the same length of time. Furthermore, a non-induced sample was included as a control for endogenous editing. Total RNAs from triplicate experiments were isolated. Libraries were generated and sequenced, and the reads were mapped to the *Drosophila* genome. Over all conditions 7,357 RNAs were detected with an expression level higher than one transcript per million (TPM). RNAs with an expression below one TPM were removed as the expression level was deemed too low. Pairwise correlation analysis of the expression level between samples and within triplicates revealed proper clustering (*Suppl Figure S2A*), indicating a high reproducibility between libraries.

To determine the extent and identity of the edited RNAs, we identified variants and used these positions to calculate the A-to-G editing frequencies for each gene (42) as above. In short, for each A-to-G variant, we divided the total amount of edits (G) by the total amount of reads for that position. Finally, the editing frequency per gene was calculated by averaging all editing frequencies per position across the same gene. In total, we found 2,844 non-edited RNAs and 4,513 edited RNAs.

We then assessed the level of endogenous editing by comparing the non-induced sample to the basal condition (induced + Schneider’s), followed by calculating and visualizing the Euclidean distance between samples. We found that 905 RNAs were endogenously edited at a low level when compared to the basal condition (**Figure 2B**). Overall, 3,608 RNAs were found to be edited by FMR1-ADARcd-V5 (*Suppl Table S1*).

**Figure 2:**
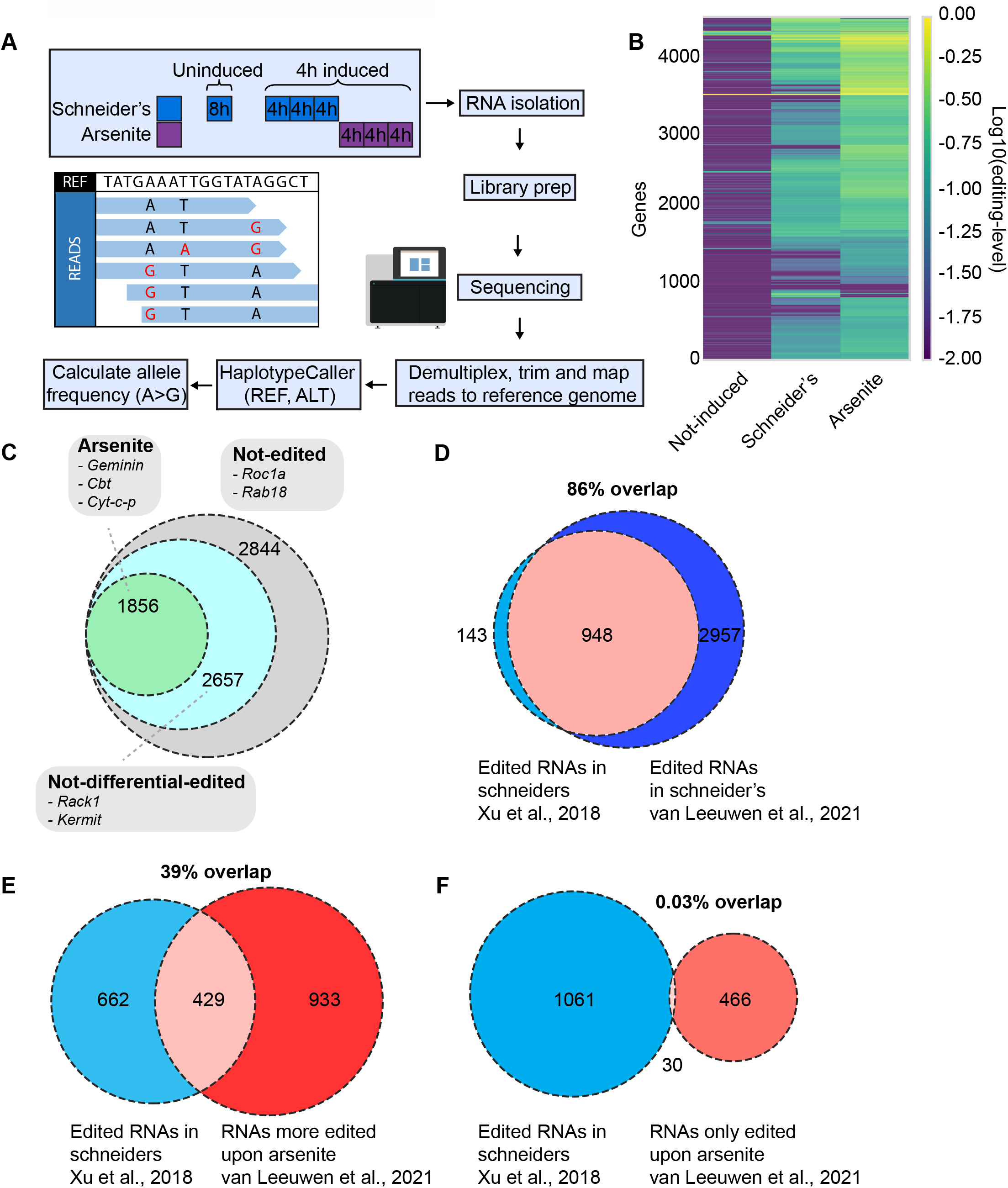
RNA editing through ADAR predicts the stress granule transcriptome in S2 cells. **A:** Schematic overview of the workflow. The expression of FMR1-ADARcd-V5 was induced for 4 h after which S2 cells are exposed to either Schneider’s or arsenite for 4 h. RNA was isolated, and libraries were generated and sequenced. Reads were processed by demultiplexing, trimming and mapping to the reference genome. The mapped reads were consequently subjected to a haplotypecaller to detect base editing events. The frequency of A>G editing events is calculated per gene and per sample. **B**: Heatmap displaying the editing level per gene and per condition. Note that there is hardly any editing in the non-induced sample (compared to schneider’s) and that editing is higher in cells treated by arsenite (as expected). **C**: Venn diagram depicting the amount of RNAs that were significantly (p<0.01) more edited upon arsenite after applying the Empirical bayes statistical test on the data set. **D**: Venn diagram depicting the percentage of RNAs that are edited in our basal condition (schneider’s) overlapping with the established 1090 FMR1 clients (28). **E**: Venn diagram depicting the percentage of RNAs that were more edited upon arsenite (when compared to basal condition) overlapping with the established FMR1 clients (28) **F**: Venn diagram depicting the percentage of RNAs that were only edited upon arsenite overlapping with the established FMR1 clients (28)

We predicted that a subset of RNAs would have higher editing levels upon arsenite stress due to their colocalization with ADARcd within stress granules. Indeed, we found 1,856 RNAs, such as *Geminin, Cyt-c-p, Red, Cbt and Khc*, that were significantly more edited upon arsenite when compared to the basal condition (p < 0.01, Empirical Bayes analysis, see material and methods) (**Figure 2B, C** and *Suppl Table S1*). Importantly, the increased editing was irrespective of the change in expression level (*Suppl Figure S2B*). The remaining 2,657 RNAs, such as *Rack1* and *Kermit*, were not differentially edited (**Figure 2C** and *Suppl Table S1*). We conclude that 1,856 RNAs are potentially recruited to stress granules upon arsenite stress.

### Potential stress granule RNAs are not all FMR1 clients

To assess whether these 1,856 RNAs are simply FMR1 clients, we compared them with the established list of FRM1 clients established by the Rosbash group (28). First, the RNAs that we found edited in basal conditions (Schneider’s) are 86% identical to the established FMR1 clients, supporting the reliability of our protocol (**Figure 2D**). Among the RNAs that are preferentially edited upon arsenite, we considered 2 groups. The first contains the RNAs that are edited at low level in basal conditions and significantly more edited in arsenite (1,360). Those are more likely to be FMR1 clients. Indeed, these overlap for 39% with the list of the FMR1 clients (**Figure 2E**, *Suppl Table S2*). The second group, however, contains the RNAs that are not edited in basal conditions (496) and only upon arsenite, and it only slightly overlaps (0.03%), with the list of the FMR1 clients (**Figure 2F**, *Suppl Table S2*). Interestingly, *Rack1* (that is a clear FMR1 client, (28)) is not recruited in stress granules (see below). This indicates that our method of identification of stress granule RNAs does not simply retrieve the clients of the RBP used for TRIBE. In our case, only 546 RNAs (out of 1,856) that are predicted to be in stress granules are clients of FMR1.

### Validation of identified stress granule transcriptome by smFISH

To validate the significant enrichment of stress specific editing of RNAs in stress granules, we used single-molecule fluorescence in situ hybridization (smFISH) to examine the localization of *Row, Geminin, Cyt-c-p, Red, Cbt* and *Khc* mRNAs in arsenite stressed wild type S2 cells. We also analyzed the localization of non-differentially edited mRNAs (*Rack1 and Kermit*) that should be depleted from stress granules (**Figure 2C**). In agreement, smFISH analysis revealed that *Row, Geminin, Cyt-c-p, Red, Cbt* and *Khc* RNA molecules strongly co-localize (54 % to 83 %) with endogenous FMR1 in stress granules **(Figure 3A-D, G**). Conversely, only 2% of *Rack1* and 29% of *Kermit* RNA overlap with stress granules (**Figure 3E-G**). This agreement within the validation set indicates that the 1,856 RNAs that we identified as being differentially edited upon arsenite treatment are indeed enriched in stress granules.

**Figure 3:**
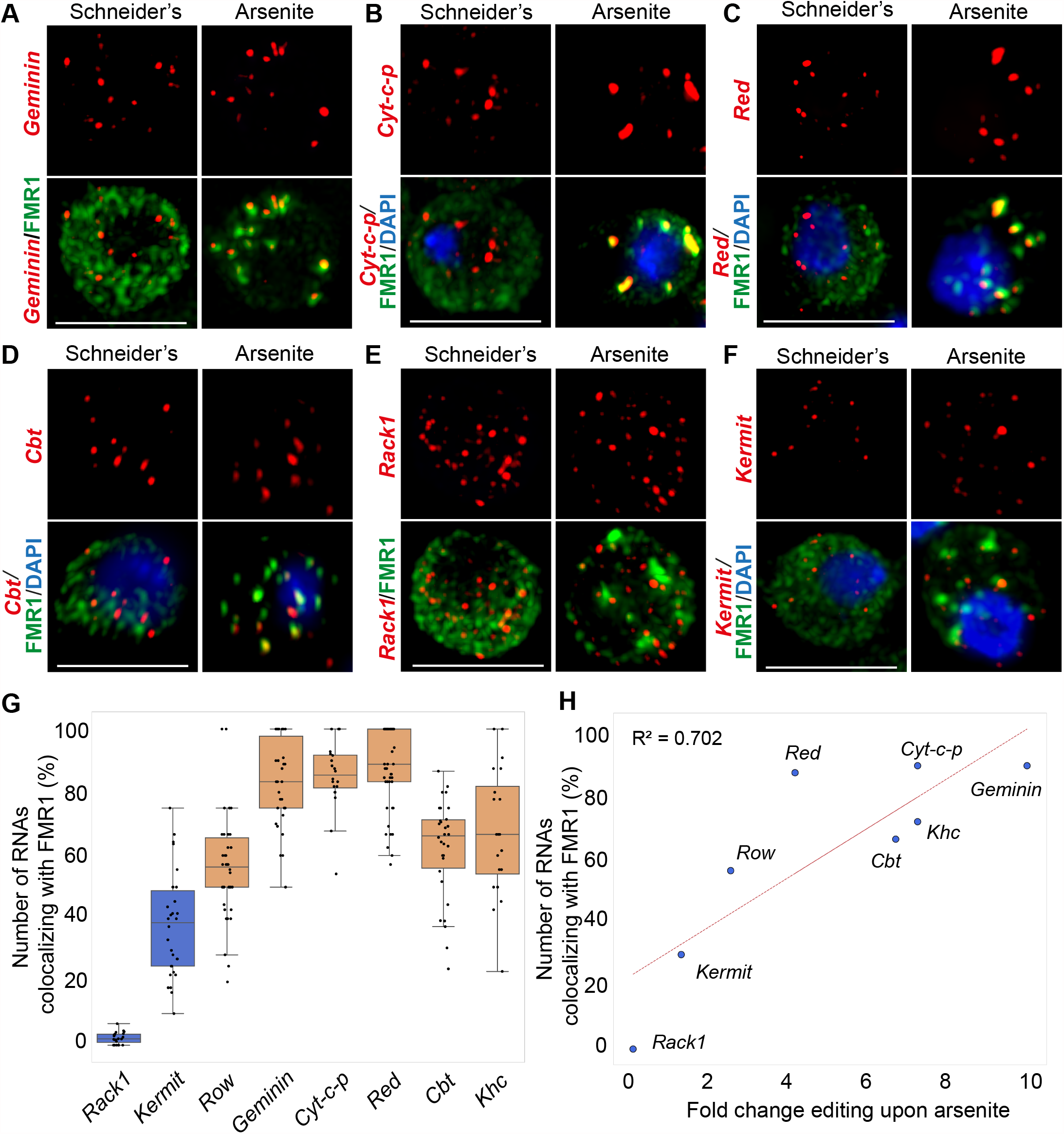
Potential stress granule RNAs identified by RNA-editing are validated by single molecule FISH. **A-D:** Visualisation of *Geminin, Cyt-c-p, Red* and *Cbt* mRNAs by smFISH, which were significantly more edited upon arsenite compared to Schneider’s(p<0.01). Note that all these mRNAs colocalize with the endogenous stress granule protein FMR1 as predicted. **E-F:** Visualisation of of *Rack1* and *Kermit* RNAs by smFISH, which were not differential edited upon arsenite compared to Schneider’s. Note that the mRNAs hardly or do not colocalize with the stress granule marker FMR1, as predicted. **G:** Quantification of the number of each indicated RNA that colocalize with FMR1 per cell. At least thirty cells were counted per tested RNA and per condition. **H:** Scatterplot displaying the number of each indicated RNA in stress granules versus the fold change editing upon arsenite compared to Schneider’s. Scale bar: 10 μm (A, E, F).

Furthermore, we set to unravel whether there is a relationship between the fold change in editing frequency of an RNA subtype (between arsenite and Schneider’s) and the fraction of RNA molecules localized in stress granules (quantified by smFISH). Indeed, we observed that an increased fold in editing reflects the localization of RNAs in stress granules (R^2^ = 0.702, **Figure 3H**). For example, *Geminin* is edited 9.7 times more in arsenite when compared to Schneider’s, and 83% of *Geminin* RNA molecules localize to stress granules. Conversely, the fold change in editing for *Rack1* is 0.7 times and the fraction of *Rack1* RNAs in stress granules is 2%. This strong correlation indicates that the higher the fold change in editing (arsenite versus basal), the most likely RNAs localize in stress granules.

Overall, these data indicate that the RNAs that were predicted to be in stress granules through RNA editing are indeed localized in stress granules.

### Arsenite-induced stress granules contain long mRNAs that encode ATP binding, transcription, cell cycle and splicing factors

We then asked whether the 1,856 stress granules RNAs that we have identified have common and specific features. First, we found that ~99.6% are mRNAs and ~0.4% are ncRNAs (**Figure 4A**). Next, we found that the stress-granule transcripts were 1.3-fold longer than the mRNAs predicted not to be in stress granules (**Figure 4B-E**). Finally, we performed gene enrichment analysis on the stress-granule mRNAs, revealing an enrichment in ATP binding (kinases, chaperones and cytoskeleton), transcription, RNA splicing, and cell cycle factors (**Figure 4F**). Together, these features suggest that stress granules may harbor and protect long RNAs that are necessary for the cells to thrive once the stress is relieved. Interestingly, these features are similar to the mammalian stress granules RNAs (21,22,24).

**Figure 4:**
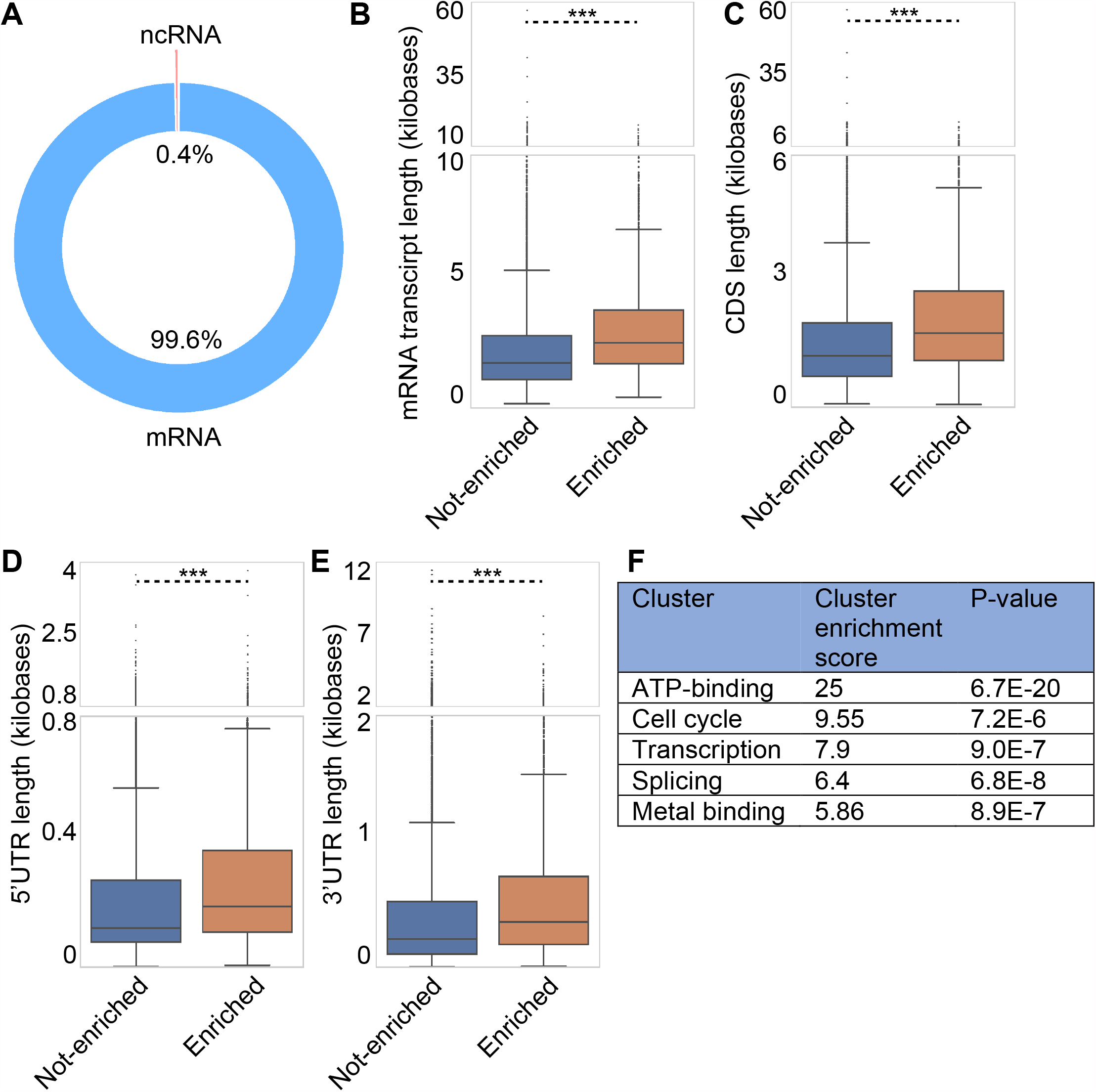
Features of the identified stress granule RNAs. **A:** Chart depicting the types of RNAs in stress granules. Note that the majority were mRNAs. **B-E:** Boxplots displaying the transcript length, CDS length, 5’UTR length and 3’UTR length of the RNAs predicted to be in stress granules (enriched) and of the RNAs that were predicted not to be localized in stress granules (not-enriched). ***p value <0.001 (student’s t-test). **F:** Gene enrichment analysis of the RNAs predicted to be localized in stress granules upon arsenite (cluster analysis using DAVID).

### Identification of stress granule RNAs in single cells

There are clear indications that the RNA content of stress granules might be heterogeneous within a cell population (22). Our own smFISH data revealed that overall, each cell does not have the same stress granule RNA enrichment as its neighbor (*Suppl Figure S3A*). The existing published techniques investigating the RNA content of stress granules require a lot of material (see introduction), which makes single cell analysis infeasible. Given that our adapted TRIBE technique does not require purification, we tested the ability of this method to identify stress granule RNAs in single cells.

The editing efficiency in bulk S2 cells was high enough to attempt detecting RNA-editing in single cells. Using the same induction and stress procedures as we previously established, we observed a significant enrichment in A-to-G editing compared to all other possible nucleotide base changes (**Figure 5A**), indicating that TRIBE can be used to detect RNA-editing in pseudobulk (combined single cells). To detect whether the editing activity changes in arsenite-stressed single cells, we calculated the editing frequency per gene and averaged this over all genes detected within a cell to determine the editing activity per cell. Interestingly, we estimated that 63 % of single cells showed a significant increase in editing activity upon arsenite treatment when compared to Schneider’s (**Figure 5B**), in agreement with our bulk RNA sequencing.

**Figure 5:**
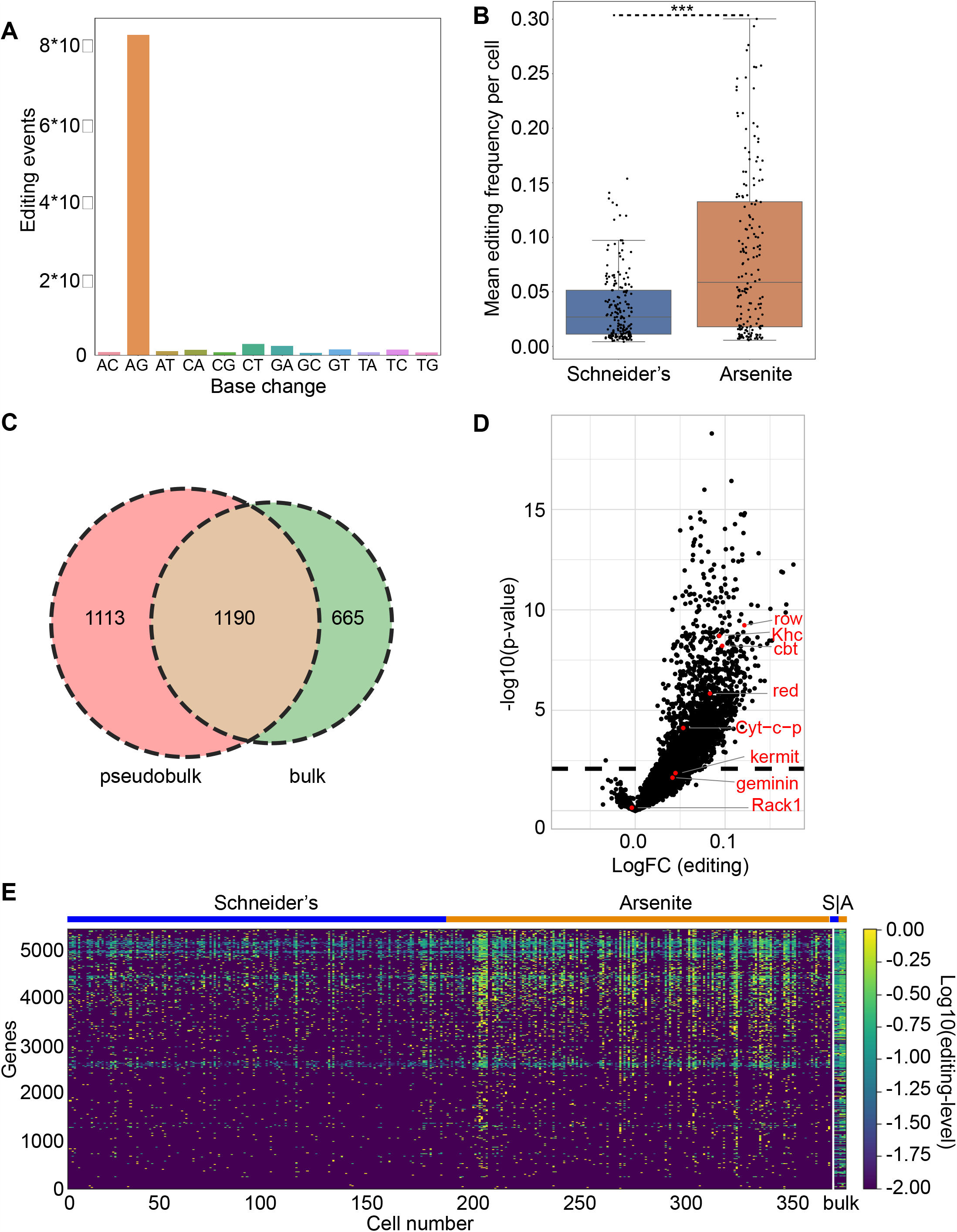
Identification of stress granule RNAs in single cells. **A:** Countplot displaying the amount of editing events per possible base change in single cells. Note that most editing events were A>G base changes. **B:** Boxplot of the mean editing frequency over all genes per single cell for each condition. ***p value <0.001. **C:** Venn diagram depicting the overlap of RNAs significantly (p<0.01) more edited upon arsenite in bulk sequencing and single cell sequencing. **D:** Volcano plot depicting the log-fold change in editing and the significancy (p<0.01) for all detected RNAs in pseudobulk. Note that most of the RNAs validated with smFISH to be enriched in stress granules were also significantly edited in pseudobulk. **E:** Heatmap of the editing level per gene per single cell. The last columns are bulk sequencing (S = Schneider’s, A = arsenite).

Performing empirical bayes statistical analysis on the pseudobulk identified 2,303 RNAs that were more edited upon arsenite. We compared those to the 1,856 RNAs found by bulk sequencing. Interestingly, 64% of the RNAs that were more edited upon arsenite in bulk are also edited in pseudobulk (**Figure 5C**). Notably, we found that most of the RNAs validated with smFISH to be enriched in stress granules were also significantly edited in pseudobulk showing the reliability of the single cell analysis (**Figure 5D**).

By comparing the editing frequency per gene between all cells, we observed a small degree of variation in the editing level between the cells, suggesting a low level of heterogeneity (**Figure 5E**).

Taken together, using our adaptation of the TRIBE method, stress granule RNAs can be reliably identified in single cells.

### Identification of stress granule RNAs in *Drosophila* neurons

We then tested whether the RNA content of stress granules may be determined in specific cell types in primary tissue and we focused on neurons of the *Drosophila* third instar larval brain. We sought to specifically express the FMR1-ADARcd-V5 fusion in *Drosophila* neurons, while allowing for precise temporal control to limit basal editing in unstressed conditions. To achieve this temporal control, we used the geneswitch/Gal4 system (43), which is a modified Gal4/UAS system (44), whereby protein expression is induced by the addition of the drug RU486. To limit the expression to neurons, we selected the pan-neuronal *elav*-geneswitch-Gal4. Thus, FMR1-ADARcd-V5 was specifically expressed in *Drosophila* neurons by bathing third instar larvae in RU486 (**Figure 6A**).

**Figure 6:**
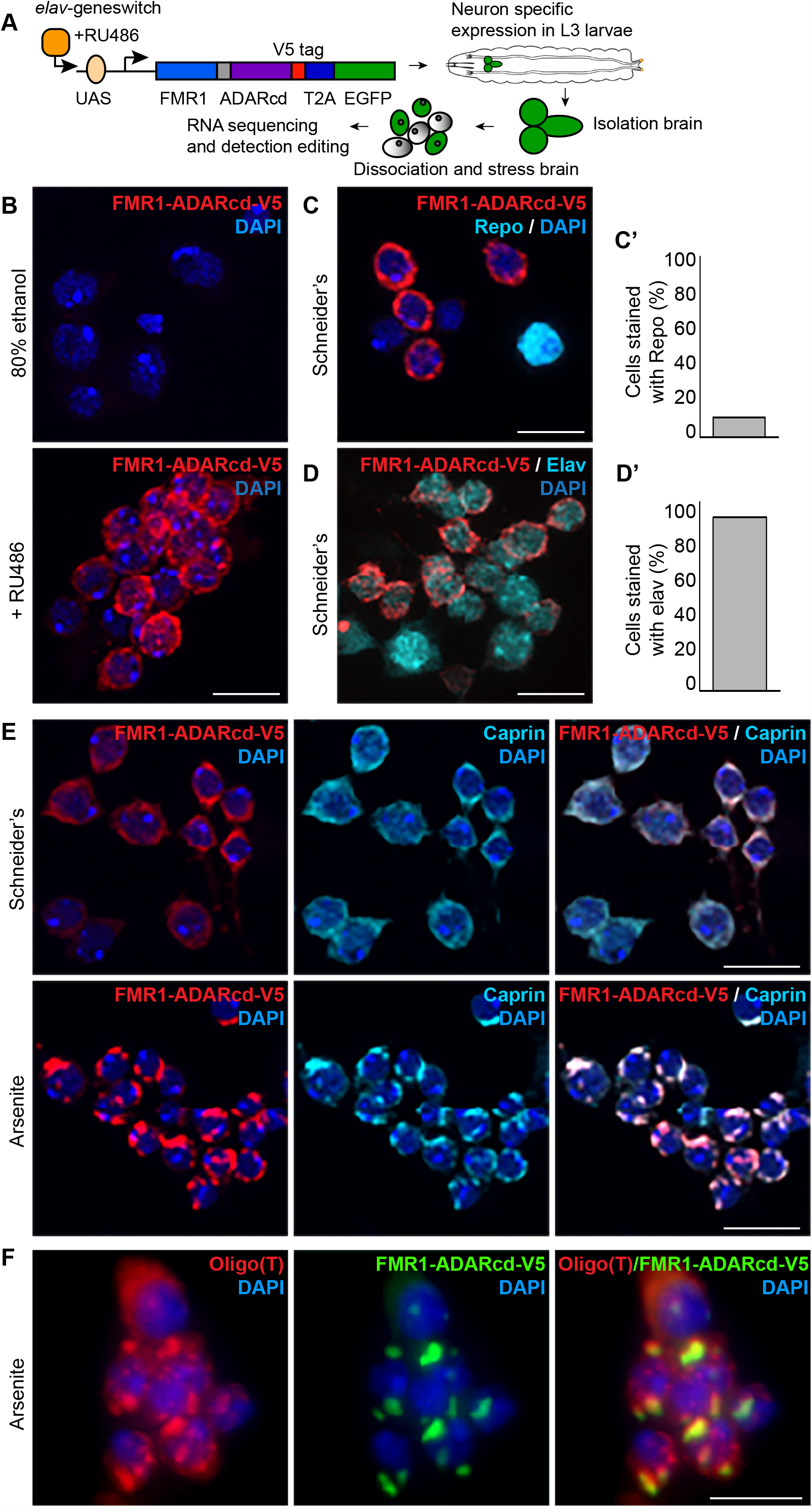
Stress granule formation in *Drosophila* neurons. **A:** Schematic overview of the workflow. *elav*-GeneSwitch-Gal4 / FMR1-ADARcd-V5 expressing third instar larvae were exposed to RU486 by larval bathing to allow the expression of FMR1-ADARcd-V5 specifically in neurons. Brains were dissected from the larvae and subsequently dissociated. The dissociated brain cells were exposed to arsenite how long and which concentration. Finally, the total RNA was isolated, sequenced, and the RNA editing analyzed. **B:** Immunofluorescence visualization of FMR1-ADARcd-V5 (red) in dissociated *Drosophila* brain cells from mock (80% ethanol) and RU486 (3 mg/ml) bathed third instar larvae. **C, C’:** Immunofluorescence visualization of FMR1-ADARcd-V5 (red) and the glial cell marker Repo (cyan) in dissociated *Drosophila* brain cells. Note that FMR1-ADARcd-V5 was not expressed in glial cells. Quantification in C’. **D, D’:** Immunofluorescence visualisation of FMR1-ADARcd-V5 (red) and the neuronal marker Elav (cyan) in dissociated *Drosophila* brain cells. Note that FMR1-ADARcd-V5 was expressed in neurons. Quantification in D’. **E, E’:** Immunofluorescence visualization of FMR1-ADARcd-V5 in dissociated *Drosophila* brain cells upon incubation in Schneider’s or in Schneider’s + 0.5mM arsenite for 4 h. Upon arsenite, FMR1-ADARcd-V5 localises in stress granules together with Caprin. Quantification of stress granule formation in E’. **F:** Overall visualization of all mRNAs in arsenite-stressed neurons by smFISH using an oligo(T) probe. Stress granules are marked with FMR1-ADARcd-V5 and contain mRNAs. Scale bar: 10 μm (B, C, D, E); 5 μm (F) Error bars: SD (C’, D’)

Dissection and dissociation of brains into isolated cells (neurons and glia) showed that 70% of them expressed FMR1-ADARcd-V5 (**Figure 6B**), whereas there was no detectable expression in the absence of the drug. To determine whether FMR1-ADARcd-V5 was solely expressed in neurons and not in glial cells, we co-stained dissociated brain cells with either the glial cell marker Repo, or with the neuronal marker elav (**Figure 6 C, D**). As expected, FMR1-ADARcd-V5 was not expressed in Repo positive cells (**Figure 6C**), while all elav positive cells (93% of total dissociated brain cells) did express FMR1-ADARcd-V5 (**Figure 6D**). This data indicates that the *elav*-GeneSwitch-Gal4 driver only allows the expression of FMR1-ADARcd-V5 in neurons and not in glial cells.

Last, we tested whether dissociated neurons form stress granules upon arsenite treatment, and whether FMR1-ADARcd-V5 is sequestered inside them, as was the case in S2 cells. Indeed, 91% of the dissociated neurons form stress granules (Caprin positive) very efficiently (i.e. they form at least one stress granule per cell) upon 4 h incubation in the presence of arsenite (**Figure 6E**). As expected, these stress granules contain mRNAs, as shown by smFISH using an oligo(dT) probe to stain the polyA tail of the mRNAs (**Figure 6F**). Overall, these data indicates that *elav*-GeneSwitch-Gal4 specifically/solely drives FMR1-ADARcd-V5 expression in neurons. Furthermore, when stressed by arsenite, these neurons form *bona fide* stress granules where FMR1-ADARcd-V5 is recruited.

### RNA editing through ADAR predicts the stress granule transcriptome of dissociated *Drosophila* neurons

We sequenced the total RNA isolated from biological triplicates of dissociated neurons from the uninduced, induced, and arsenite-stressed conditions. Across all conditions, we detected the expression of 8,505 genes, almost 50 % of which were edited (4,137 of 8,505; *Suppl Table S3*).

By comparing the editing level per gene across the conditions, we found that there is hardly any editing when FMR1-ADARcd-V5 is not expressed, indicating that the endogenous A-to-G editing in neurons is very low **(Figure 7A)**. However, when FMR1-ADARcd-V5 was expressed, a clear increase in editing was observed, indicating that ADARcd is active and able to edit RNA in neurons.

**Figure 7:**
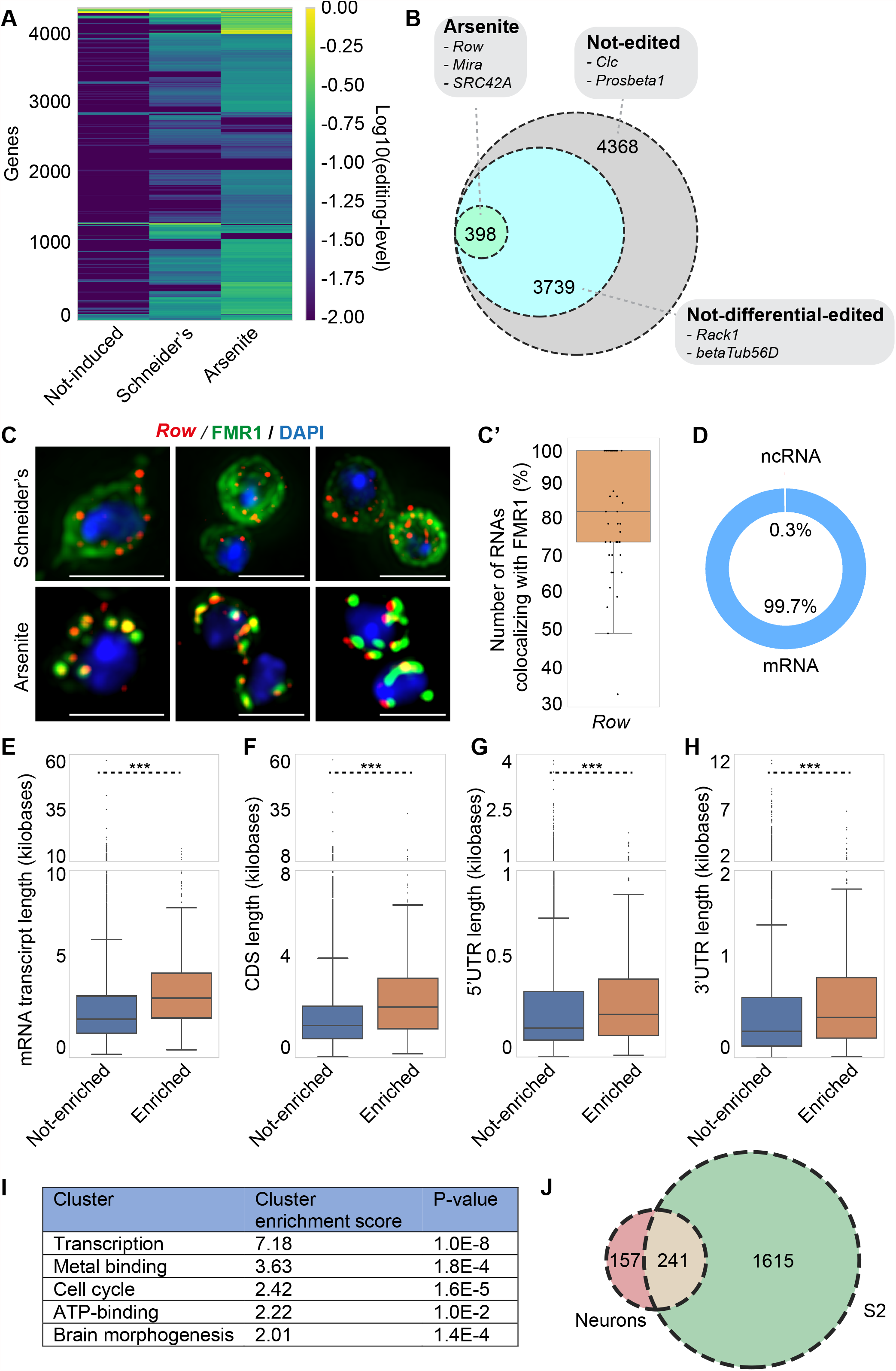
RNA editing through ADAR predicts the stress granule transcriptome of dissociated *Drosophila* neurons. **A:** Heatmap displaying the editing level per gene and per condition. Note that there was hardly any editing in the non-induced sample when compared to Schneider’s, and that upon arsenite, the level of editing increases further. **B:** Venn diagram depicting the amount of RNAs that were significantly (p<0.01) more edited upon arsenite after applying the Empirical bayes statistical test on the data set. **C:** Visualization of the *Row* mRNA by smFISH in wild-type neurons, which was significantly (p<0.01) more edited upon arsenite compared to Schneider’s in neurons. Stress granules are marked with endogenous FMR1. Note, that *Row* is localized in stress granules. Quantification of the number of *Row* RNAs co-localizing with FMR1 (stress granules) in C’. **D:** Chart depicting the types of RNAs in stress granules. Note that the majority were mRNAs. **E-H:** Boxplots displaying the transcript length, CDS length, 5’UTR length and 3’UTR length of the RNAs predicted to be in stress granules (enriched) and of the RNAs that were predicted not to be localized in stress granules (not-enriched) in neurons. ***p value <0.001 (student’s t-test). **I:** Gene enrichment analysis of the RNAs predicted to be localized in stress granules upon arsenite in neurons (cluster analysis using DAVID). **J:** Venn diagram depicting the overlap of RNAs significantly (p<0.01) more edited upon arsenite in S2 cells and in *Drosophila* neurons. Scale bar: 3 μm (C)

Importantly, as for S2 cells, many RNAs were more edited upon arsenite compared to Schneider’s and certain RNAs are only edited upon arsenite. We identified 398 RNAs, such as *Row, Mira*, and *SRC42A* that were significantly more edited upon arsenite stress (p < 0.01, Emperical bayes test) (**Figure 7B**). The remainder 3,739 RNAs, such as *Rack1* were not-differentially edited. TRIBE therefore predicts that 398 RNAs are specifically recruited to neuronal stress granules upon arsenite treatment.

Following up on these predictions we used smFISH to validate that the RNA transcript *Row* is localized in arsenite-induced stress granules in the neurons of wild-type flies as predicted (**Figure 7C, C’**). These data indicate that, similarly to S2 cells, transcripts that were predicted to be in stress granules through RNA editing are indeed localized in stress granules.

### Arsenite-induced stress granules in S2 cells and neurons share the same RNAs

As in S2 cells, we found the majority (~99.7%) of RNAs predicted to be recruited to neuronal stress granules are coding mRNAs and ~0.3% are ncRNAs (**Figure 7D**). The analysis of their mean mRNA length indicated that they are on average longer than mRNA not recruited in stress granules (**Figure 7E-H**). This suggests that, as for S2 cells, the length of mRNA might be an important feature for the recruitment to stress granules. Last, gene enrichment analysis revealed that, as in S2 cells, the predicted stress granule RNAs show an enrichment for genes encoding ATP binding, transcription, cell cycle factors (**Figure 7I**).

Performing a parallel analysis in Drosophila S2 cells and neurons allowed us to assess to which extent the RNA content of stress granules is shared. Out of the 398 RNAs predicted in neuronal stress granules, 241 (61%) were also predicted to be in stress granules in S2 cells (**Figure 7J**), suggesting a large overlap. The remaining 39% are therefore neuron specific and encode neuron related factors/proteins, such as *Mira*.

## DISCUSSION

Here, using our adaptation of RNA editing via TRIBE, we show that the RNA content of stress granules can be reliably identified not only in bulk cells but also in single cells (Drosophila S2 cells) and tissues *(Drosophila* neurons).

### Identification of stress granule transcriptome in single cells

In bulk S2 cells, we identified 1,856 potential stress granules RNAs, 6 of them validated by smFISH. The results obtained with our method compare well with those published for mammalian cells as described in the introduction (21,22,24). First, the number of stress granule RNAs we identified is similar to the number found for mammalian cells (1,800 RNAs in (22) and 1,400 RNAs in (21), strengthening the notion that stress granules harbor a large variety of RNAs. Second, as in mammalian (21,22,24), and yeast cells (22), 99% of the stress granule RNAs in S2 cells are mRNAs that are longer (here 30%) than their non-recruited counterparts. Longer mRNAs may potentially have more binding sites for RBPs that are proposed to be major driving factors in stress granules partitioning (15–17,25,45–47). They may also extend beyond the stress granule physical boundaries where they might interact with, and pull, cytoplasmic RNAs within stress granules (48). Indeed, RNA-RNA interactions are proposed to contribute to stress granule assembly (49), and this is reinforced by the notion that RNAs themselves can phase separate (20). Third, the gene ontology analysis of the S2 cell stress granule mRNAs shows enrichment in ATP binding, transcription, cell cycle and RNA splicing factors that also compare well the stress granule mRNA complement in NIH3T3 cells (21). These similarities indicates that a common core of stress granule RNAs might be conserved across evolution.

Importantly, when compared to other published techniques, the RNA-editing based method we developed has the major advantage to not require isolation of stress granules. Our method preserves cell integrity and requires only a small amount of edited RNA, thus allowing analysis in single cells for the first time. Importantly, we found an overlap of 64% of the edited RNAs between the pseudobulk and the bulk analysis, showing the reliability of the approach. The analysis of these editing events did not reveal a heterogeneity between cells, either those kept in basal conditions, or those that were arsenite stressed. However, heterogeneity in cultured cells in S2 cells is not necessarily expected. Our method could therefore be used for the detection stress granules RNAs in rare cells *in vivo*, either forming a small tissue, or disseminated into larger tissues.

### Drosophila Neurons

In this regard, using genetic tools to control spatial and temporal expression of ADARcd, we show that our method can be adapted to specifically identify the arsenite triggered stress granule transcriptome in *Drosophila* neurons. Neuronal stress granules contain less RNAs (~5% of the expressed transcriptome) than S2 cells (~25%). Ruling out technical differences in ribosome removal and sequencing, these differences might be due to the small size of neurons (3μm in diameter) when compared to S2 cells (10μm), a lower number of stress granules formed in neurons (~3/cell) than in S2 cells (~15/cell), and the lower editing efficiency in neurons. Despite these differences, 398 RNAs predicted to be in neuronal stress granules have been identified (and validated one (*Row*) by smFISH) with features similar as those of S2 cell stress granules, i.e. mainly longer mRNAs displaying a similar enrichment for ATP binding, transcription and cell cycle factors. The fact that cell cycle factors are found in neuronal stress granules might somewhat be surprising, as mammalian adult neurons are not known to be dividing, and they only represent 10-20% of the cells in the human brain (50,51). However, neurons represent 90% in the *Drosophila* brain cells, especially in larval brain that contain many neuroblasts, the neuron precursors (also expressing elav) that are still dividing (52,53).

The identification of the stress granule transcriptome in tissues and in specific cells within a given tissue opens the possibility for further analysis in other organs. Furthermore, this technique could be extended to mammalian cells and tissues through the use of TET-on inducible system and specific conditional CRE promotors to ensure tissue and time specificity. This could also be easily followed by single cell analysis, to not only investigate the stress granule RNA content in specific cell type, but also to address their heterogeneity of their response.

### Limitations

However, this successful proof of principle presents several limitations. First, cells and tissues need to express ADARcd through transfection or generation of transgenic animals. Second, one of the requirements is to limit the basal editing (in the absence of stress) by controlling the timing and temporal (in *in vivo*) expression of ADARcd. In *Drosophila* cells and tissues, we have solved this issue by using the metallothionein inducible promoter and the geneswitch system. Photoactivatable (Light oxygen voltage) FRM1-ADARcd would also be envisioned for specific activation in stress granules, thus to restrict even more the background (54). Third, certain RNAs may be covered with RNA-binding proteins therefore blocking the editing sites. Fourth, once edited, a given RNA molecule might leave the stress granule and return to the cytoplasm or to P-bodies (55,56), leading to the identification of RNAs that are only transiently residing in stress granules. Last, ADARcd was only tagged to one RBP (FMR1), thus potentially biasing our analysis toward FMR1 clients. However, we clearly showed that it was not the case and that 60% of the stress granules RNAs we identified are not FMR1 clients.

### Single molecule FISH

Out of the potential stress granule RNAs that we have identified, 6 have been validated by smFISH, plus 2 that are predicted not to be. An extended validation could be performed using multiplex smFISH (57). This could reveal whether stress granules are populated with all RNAs identified, or a subset of them, and in which ratio. If such a heterogenous recruitment is observed, understanding what governs this heterogeneity could reveal important principles governing stress granule formation. Our smFISH in S2 cells has begun to reveal such a heterogeneity (*Suppl Figure S3A-B*). The variety observed per cell and per stress granules may reflect the large variability in their RNA recruitment (**Figure 3C**).

### Why are RNAs recruited to stress granules?

One of the large questions in the stress granule field is why mRNAs are recruited in stress granules. Besides their structural role that has been proposed recently (20,49), one general assumption is that stress granules form upon stress to protect RNA from degradation after translation stalls and that

RNAs are no longer covered in ribosomes, thus allowing binding to stress granule RBPs. When stress is relieved, these protected and capped mRNAs could be immediately translated enabling the cells to thrive again and gain a fitness advantage. Indeed, stress granule formation has been shown to allow cell survival upon stress and thriving upon stress relief (58–60). The enrichment in RNAs encoding ATP binding, cell cycle and transcription factors may be consistent with this notion. The enrichment of RNAs encoding splicing factors might also be somehow related to the observation that the inhibition of RNA splicing leads to stress granule formation (61)

However, contrary to the finding that stress granules do not contain fully assembled ribosomes (4,12), single-molecule imaging has revealed that the translation machinery can be activated in stress granules (62). This suggests that although general translation is inhibited, stress granules could be a hub for translating specific RNAs, perhaps related to coping with stress.

As a proof of principle, the development of this technique opens avenues to refine the specific core of common stress granule RNAs upon different stress and understand their role and their features. It is the first method to identify stress granule RNAs in single cells and in tissues, and could be particularly important to reveal heterogeneity between cells, within primary cells or tissues in their response to different stress. Importantly, this method could also allow the identification the RNA content of other RNA based phase separated assemblies, whether formed upon stress or existing in basal conditions, such as neuronal granules (63,64), posterior granules in Drosophila oocyte (65,66) and P granules in C.elegans (67,68), whose RNA content has so far been difficult to isolate and study.

## DATA AND SOFTWARE AVAILABILITY

Raw sequencing data have been made available in the Gene Expression Omnibus under the accession number GSE175782. All scripts to process raw data are available at https://github.com/mvanins/stress_granule_RNA_manuscript.

## ACKNOWLEDGEMENTS

We would like to thank Geert Kops and Jacques Bothma for critically reading the manuscript. We would also like to thank Anko de Graaff of the Hubrecht Imaging Facility for helping with the imaging and Buys de Barbanson for helping with data analysis.

## LEGENDS FOR SUPPLEMENTARY FIGURES

**Suppl Figure S1:**
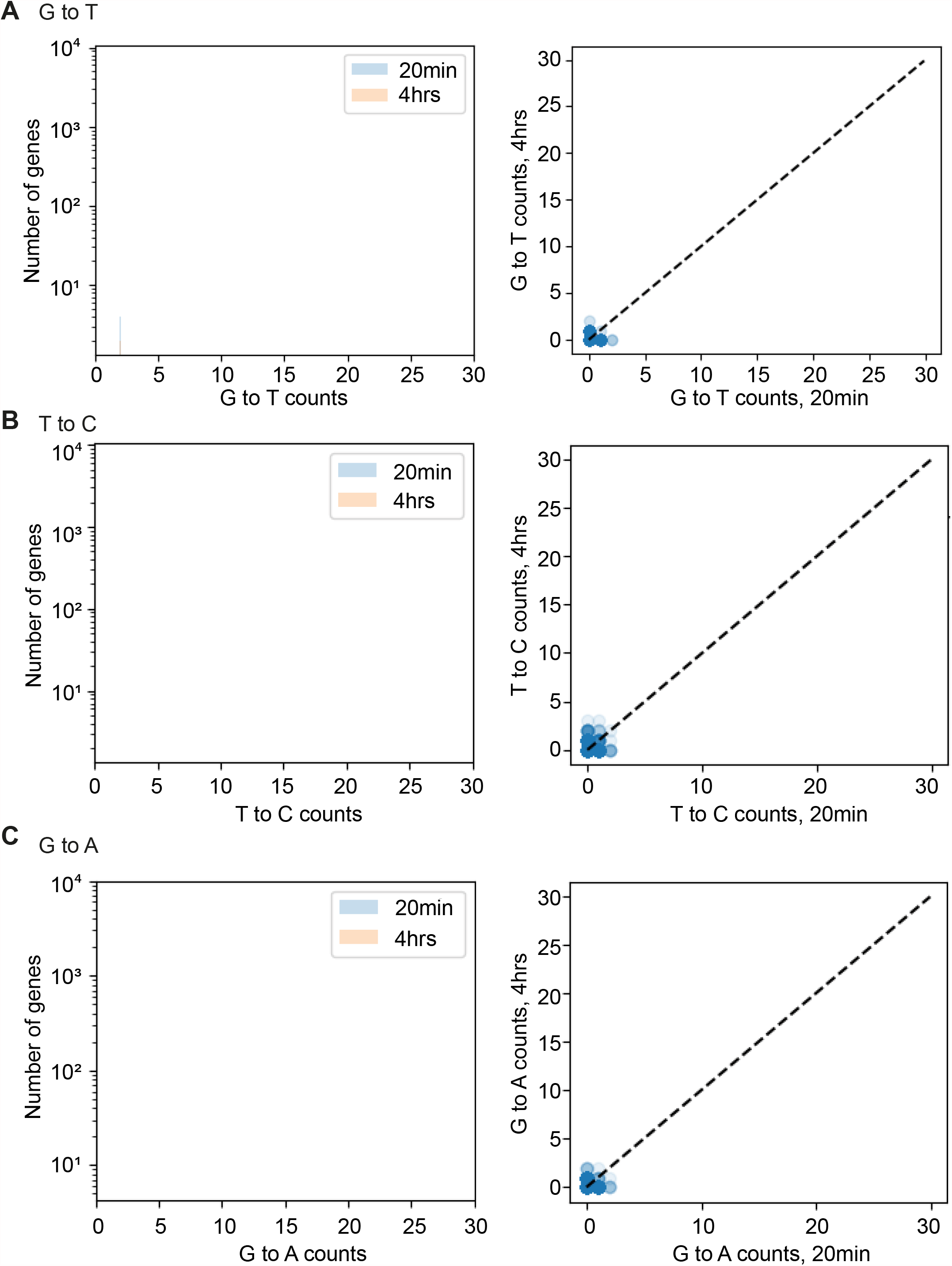
Frequency of other possible base changes. **A-C:** Graph displaying the frequency of G>T, T>C or G>A editing events in S2 cells in which the expression of FMR1-ADARcd-V5 was induced for 20 min or 4 h. Note that there were no editing events for these combinations.

**Suppl Figure S2:**
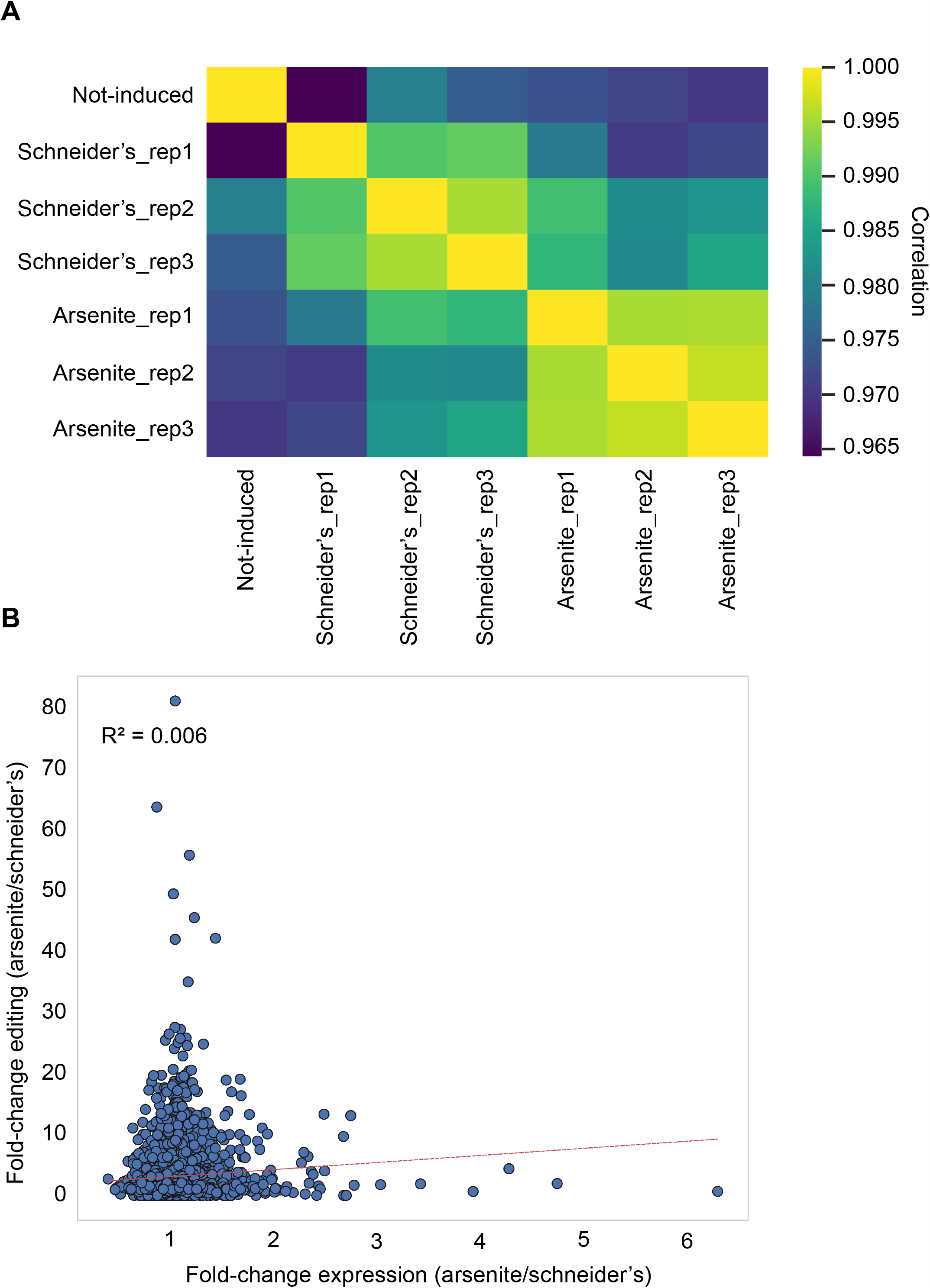
Correlation RNA expression levels and comparing expression level vs fold-change editing. **A:** Heatmap displaying the RNA expression level correlation between the triplicates of S2 cells. **B:** Scatter plot comparing the fold-change in expression level and fold-change in editing between arsenite and Schneider’s. Note that some RNAs that do not change in expression (1) are edited heavily upon arsenite.

**Suppl Figure S3:**
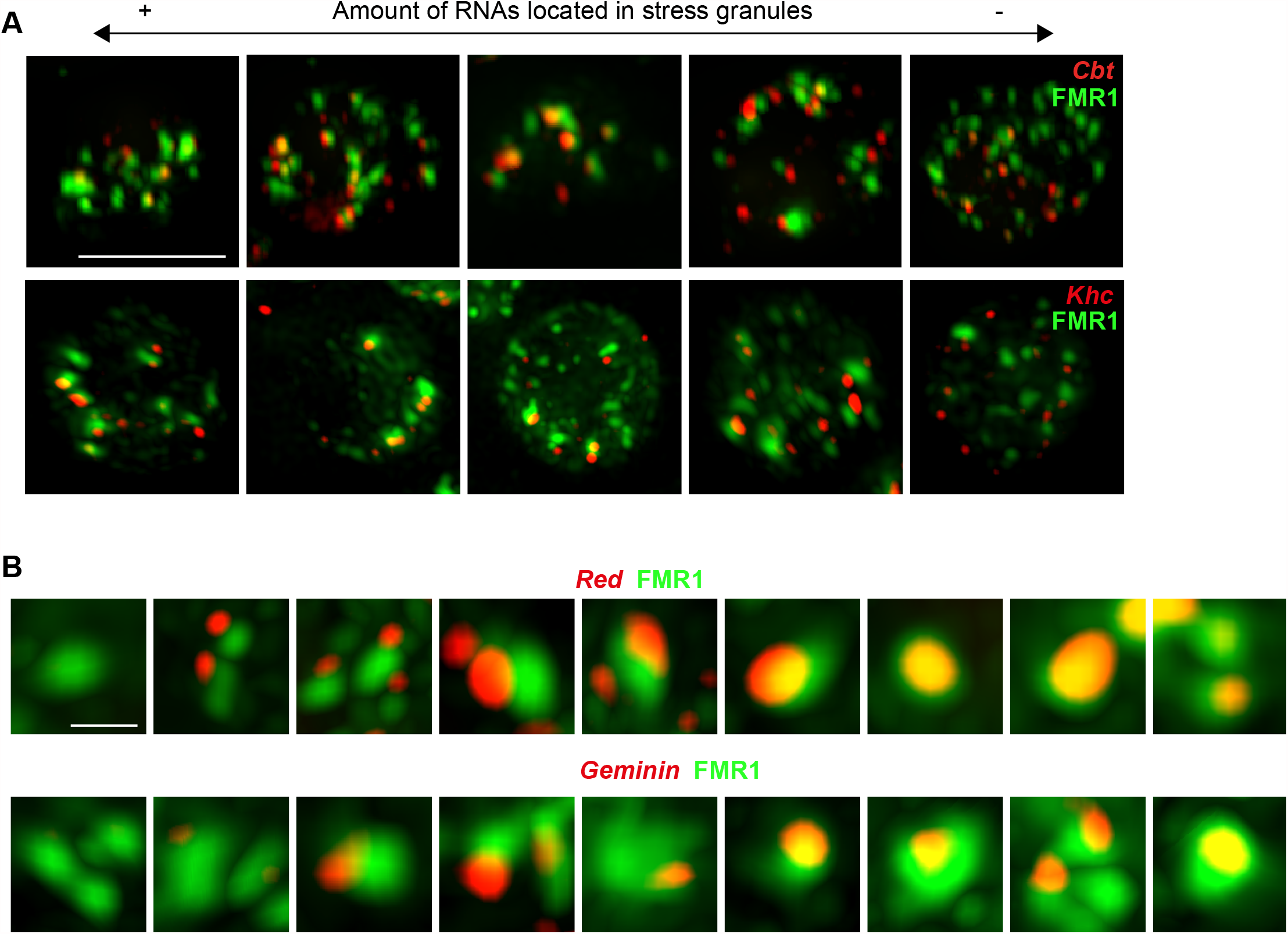
smFISH depicting the heterogeneity between cells and stress granules. **A:** Visualisation of *Cbt* and *Khc* RNAs by smFISH in S2 cells (red) showing the heterogeneity in the amount of RNA molecules located in stress granules (FMR1). Cells on the far left have more RNA molecules in stress granules that cells on the far right. **B:** Visualisation of *Red* and *Geminin* mRNAs in smFISH showing that stress granules (marked by FMR1, red) can be heterogenous in RNA content. Stress granules on the far left do not contain the tested RNA molecules, while stress granules on the far right contain many of tested RNA molecules. Scale bar: 10 μm (A), 1 μm (B).

*Supplementary Table S1*: RNA editing S2 cells

*Supplementary Table S2*: Overlap with established FMR1 clients (28)

*Supplementary Table S3*: RNA editing neurons

*Supplementary Table S4*: oligos

## REFERENCES

1. Hyman, A.A., Weber, C.A. and Julicher, F. (2014) Liquid-liquid phase separation in biology. Annual review of cell and developmental biology, 30, 39–58.

2. Gomes, E. and Shorter, J. (2018) The molecular language of membraneless organelles. Journal of Biological Chemistry, pii: jbc.TM118.001192. doi:.

3. van Leeuwen, W. and Rabouille, C. (2019) Cellular stress leads to the formation of membraneless stress assemblies in eukaryotic cells. Traffic, 0.

4. Anderson, P. and Kedersha, N. (2002) Stressful initiations. Journal of Cell Science, 115, 3227–3234.

5. Aulas, A., Fay, M.M., Lyons, S.M., Achorn, C.A., Kedersha, N., Anderson, P. and Ivanov, P. (2017) Stress-specific differences in assembly and composition of stress granules and related foci. J Cell Sci, 130, 927–937.

6. Anderson, P. and Kedersha, N. (2008) Stress granules: the Tao of RNA triage. Trends in biochemical sciences, 33, 141–150.

7. Anderson, P. and Kedersha, N. (2009) Stress granules. Current Biology, 19, R397–398.

8. Jain, S., Wheeler, J.R., Walters, R.W., Agrawal, A., Barsic, A. and Parker, R. (2016) ATPase-Modulated Stress Granules Contain a Diverse Proteome and Substructure. Cell, 164, 487–498.

9. Youn, J.Y., Dyakov, B.J.A., Zhang, J., Knight, J.D.R., Vernon, R.M., Forman-Kay, J.D. and Gingras, A.C. (2019) Properties of Stress Granule and P-Body Proteomes. MOl Cell, 76, 286–294.

10. Markmiller, S., Soltanieh, S., Server, K.L., Mak, R., Jin, W., Fang, M.Y., Luo, E.C., Krach, F., Yang, D., Sen, A. et al.. (2019) Context-Dependent and Disease-Specific Diversity in Protein Interactions within Stress Granules. Cell, 172, 590–604.

11. Tourriere, H., Chebli, K., Zekri, L., Courselaud, B., Blanchard, J.M., Bertrand, E. and Tazi, J. (2003) The RasGAP-associated endoribonuclease G3BP assembles stress granules. J Cell Biol, 160, 823–831.

12. Kedersha, N.L., Gupta, M., Li, W., Miller, I. and Anderson, P. (1999) RNA-binding proteins TIA-1 and TIAR link the phosphorylation of eIF-2 alpha to the assembly of mammalian stress granules. J Cell Biol, 147, 1431–1442.

13. Solomon, S., Xu, Y., Wang, B., David, M.D., Schubert, P., Kennedy, D. and Schrader, J.W. (2007) Distinct structural features of caprin-1 mediate its interaction with G3BP-1 and its induction of phosphorylation of eukaryotic translation initiation factor 2alpha, entry to cytoplasmic stress granules, and selective interaction with a subset of mRNAs. Mol Cell Biol, 27, 2324–2342.

14. Kedersha, N., Cho, M.R., Li, W., Yacono, P.W., Chen, S., Gilks, N., Golan, D.E. and Anderson, P. (2000) Dynamic shuttling of TIA-1 accompanies the recruitment of mRNA to mammalian stress granules. J Cell Biol, 151, 1257–1268.

15. Patel, A., Lee, H.O., Jawerth, L., Maharana, S., Jahnel, M., Hein, M.Y., Stoynov, S., Mahamid, J., Saha, S., Franzmann, T.M. et al.. (2015) A Liquid-to-Solid Phase Transition of the ALS Protein FUS Accelerated by Disease Mutation. Cell, 162, 1066–1077.

16. Murray, D.T., Kato, M., Lin, Y., Thurber, K.R., Hung, I., McKnight, S.L. and Tycko, R. (2017) Structure of FUS Protein Fibrils and Its Relevance to Self-Assembly and Phase Separation of Low-Complexity Domains. Cell, 171, 615–627.

17. Molliex, A., Temirov, J., Lee, J., Coughlin, M., Kanagaraj, A.P., Kim, H.J., Mittag, T. and Taylor, J.P. (2015) Phase Separation by Low Complexity Domains Promotes Stress Granule Assembly and Drives Pathological Fibrillization. Cell, 163, 123–133.

18. Zhang, H., Elbaum-Garfinkle, S., Langdon, E.M., Taylor, N., Occhipinti, P., Bridges, A.A., Brangwynne, C.P. and Gladfelter, A.S. (2015) RNA Controls PolyQ Protein Phase Transitions. Mol Cell, 60, 220–230.

19. Van Treeck, B., Protter, D.S.W., Matheny, T., Khong, A., Link, C.D. and Parker, R. (2018) RNA self-assembly contributes to stress granule formation and defining the stress granule transcriptome. Proc Natl Acad Sci U S A, 115, 2734–2739.

20. Langdon, E.M., Qiu, Y. N.A. G., McLaughlin, G.A., Weidmann, C.A., Gerbich, T.M., Smith, J.A., Crutchley, J.M., Termini, C.M., Weeks, K.M. et al.. (2018) mRNA structure determines specificity of a polyQ-driven phase separation. Science, 360, 922.

21. Namkoong, S., Ho, A., Woo, Y.M., Kwak, H. and Lee, J.H. (2018) Systematic Characterization of Stress-Induced RNA Granulation. Molecular Cell, 70, 175-187.e178.

22. Khong, A., Matheny, T., Jain, S., Mitchell, S.F., Wheeler, J.R. and Parker, R. (2017) The Stress Granule Transcriptome Reveals Principles of mRNA Accumulation in Stress Granules. Mol Cell, 68, 808–820.

23. Anders, M., Chelysheva, I., Goebel, I., Trenkner, T., Zhou, J., Mao, Y., Verzini, S., Qian, S.B. and Ignatova, Z. (2018) Dynamic m(6)A methylation facilitates mRNA triaging to stress granules. Life Sci Alliance, 3, e201800113.

24. Padrón, A., Iwasaki, S. and Ingolia, N.T. (2019) Proximity RNA Labeling by APEX-Seq Reveals the Organization of Translation Initiation Complexes and Repressive RNA Granules. Mol Cell, 75, 875–887.

25. Matheny, T., Van Treeck, B., Huynh, T.N. and Parker, R. (2021) RNA partitioning into stress granules is based on the summation of multiple interactions. RNA, 27, 174–189.

26. Wheeler, J.R., Matheny, T., Jain, S., Abrisch, R. and Parker, R. (2016) Distinct stages in stress granule assembly and disassembly. eLife, 5, e18413.

27. McMahon, A.C., Rahman, R., Jin, H., Shen, J.L., Fieldsend, A., Luo, W. and Rosbash, M. (2016) TRIBE: Hijacking an RNA-Editing Enzyme to Identify Cell-Specific Targets of RNA-Binding Proteins. Cell, 165, 742–753.

28. Xu, W., Rahman, R. and Rosbash, M. (2017) Mechanistic implications of enhanced editing by a HyperTRIBE RNA-binding protein. RNA, 24, 173–182.

29. Aguilera-Gomez, A., Zacharogianni, M., van Oorschot, M.M., Genau, H., Grond, R., Veenendaal, T., Sinsimer, K.S., Gavis, E.R., Behrends, C. and Rabouille, C. (2017) Phospho-Rasputin Stabilization by Sec16 Is Required for Stress Granule Formation upon Amino Acid Starvation. Cell reports, 20, 935–948.

30. Rinaldini, L.M. (1959) An improved method for the isolation and quantitative cultivation of embryonic cells. Exp Cell Res, 16, 477–505.

31. Reich, J. and Papoulas, O. (2012) Caprin controls follicle stem cell fate in the Drosophila ovary. PLoS One, 7, e35365.

32. Gaspar, I., Wippich, F. and Ephrussi, A. (2017) Enzymatic production of single-molecule FISH and RNA capture probes. RNA, 23, 1582–1591.

33. Sage, D., Donati, L., Soulez, F., Fortun, D., Schmit, G., Seitz, A., Guiet, R., Vonesch, C. and Unser, M. (2017) DeconvolutionLab2: An open-source software for deconvolution microscopy. Methods, 115, 28–41.

34. Raj, A., van den Bogaard, P., Rifkin, S.A., van Oudenaarden, A. and Tyagi, S.C. (2008) Imaging individual mRNA molecules using multiple singly labeled probes. Nature Methods, 5, 877–879.

35. Muraro, M.J., Dharmadhikari, G., Grün, D., Groen, N., Dielen, T., Jansen, E., van Gurp, L., Engelse, M.A., Carlotti, F., de Koning, E.J. et al.. (2016) A Single-Cell Transcriptome Atlas of the Human Pancreas. Cell Syst., 3, 385–394.

36. Lowe, T.M. and Chan, P.P. (2016) tRNAscan-SE On-line: integrating search and context for analysis of transfer RNA genes. Nucleic Acids Research, 44, W54–W57.

37. Martin, M.A. (2011) Cutadapt removes adapter sequences from high-throughput sequencing reads. EMBnet.journal; Vol 17, No 1: Next Generation Sequencing Data AnalysisDO - 10.14806/ej.17.1.200.

38. Li, H. and Durbin, R. (2009) Fast and accurate short read alignment with Burrows–Wheeler transform. Bioinformatics, 25, 1754–1760.

39. Kaminow, B., Yunusov, D. and Dobin, A. (2021) STARsolo: accurate, fast and versatile mapping/quantification of single-cell and single-nucleus RNA-seq data. bioRxiv, 2021.2005.2005.442755.

40. Smith, T., Heger, A. and Sudbery, I. (2017) UMI-tools: modeling sequencing errors in Unique Molecular Identifiers to improve quantification accuracy. Genome Research, 27, 491–499.

41. DePristo, M.A., Banks, E., Poplin, R., Garimella, K.V., Maguire, J.R., Hartl, C., Philippakis, A.A., del Angel, G., Rivas, M.A., Hanna, M. et al.. (2011) A framework for variation discovery and genotyping using next-generation DNA sequencing data. Nature Genetics, 43, 491–498.

42. McKenna, A., Hanna, M., Banks, E., Sivachenko, A., Cibulskis, K., Kernytsky, A., Garimella, K., Altshuler, D., Gabriel, S., Daly, M. et al.. (2010) The Genome Analysis Toolkit: a MapReduce framework for analyzing next-generation DNA sequencing data. Genome Res, 20, 1297–1303.

43. Osterwalder, T. S. Y.K. White, B.H. and Keshishian, H. (2001) A conditional tissue-specific transgene expression system using inducible GAL4. Proceedings of the National Academy of Sciences of the United States of America, 98, 12596–12601.

44. Phelps, C.B. and Brand, A.H. (1998) Ectopic gene expression in Drosophila using GAL4 system. Methods, 14, 367–379.

45. Maharana, S., Wang, J., Papadopoulos, D.K., Richter, D., Pozniakovsky, A., Poser, I., Bickle, M., Rizk, S., Guillén-Boixet, J., Franzmann, T.M. et al.. (2018) RNA buffers the phase separation behavior of prion-like RNA binding proteins. Science, 360, 918.

46. Banani, S.F., Lee, H.O., Hyman, A.A. and Rosen, M.K. (2017) Biomolecular condensates: organizers of cellular biochemistry. Nat Rev Mol Cell Biol, 18, 285–298.

47. Jain, A. and Vale, R.D. (2017) RNA phase transitions in repeat expansion disorders. Nature, 546, 243–247.

48. Moon, S.L. and Parker, R. (2018) Analysis of eIF2B bodies and their relationships with stress granules and P-bodies. Scientific Reports, 8, 12264.

49. Van Treeck, B. and Parker, R. (2018) Emerging Roles for Intermolecular RNA-RNA Interactions in RNP Assemblies. Cell, 174, 791–802.

50. Bittern, J., Pogodalla, N., Ohm, H., Brüser, L., Kottmeier, R., Schirmeier, S. and Klämbt, C. (2020) Neuron-glia interaction in the Drosophila nervous system. LID - 10.1002/dneu.22737 [doi]. Dev Neurobiol.

51. Herculano-Houzel, S. (2014) The glia/neuron ratio: how it varies uniformly across brain structures and species and what that means for brain physiology and evolution. Glia, 62, 1377–1391.

52. Homem, C.C. and Knoblich, J.A. (2012) Drosophila neuroblasts: a model for stem cell biology. Development, 139, 4297–4310.

53. Harding, K. and White, K. (2019) Drosophila as a Model for Developmental Biology: Stem Cell-Fate Decisions in the Developing Nervous System.. Developmental biology, 456, 17–24.

54. Wu, Y.I. D. F., Lungu, O.I., Jaehrig, A. I. S., Kuhlman, B. and Hahn, K.M. (2009) A genetically encoded photoactivatable Rac controls the motility of living cells. Nature, 461, 104–108.

55. Zhang, J. K. O. T. T. and Funatsu, T. (2011) Dynamic association-dissociation and harboring of endogenous mRNAs in stress granules. J. Cell Sci., 124, 4087–4095.

56. Mollet, S., Cougot, N., Wilczynska, A., Dautry, F., Kress, M., Bertrand, E. and Weil, D. (2008) Translationally Repressed mRNA Transiently Cycles through Stress Granules during Stress. Molecular Biology of the Cell, 19, 4469–4479.

57. Maynard, K.R., Tippani, M., Takahashi, Y., Phan, B.N., Hyde, T.M., Jaffe, A.E. and Martinowich, K. (2020) dotdotdot: an automated approach to quantify multiplex single molecule fluorescent in situ hybridization (smFISH) images in complex tissues. Nucleic Acids Res., 48.

58. Jevtov, I., Zacharogianni, M., van Oorschot, M.M., van Zadelhoff, G., Aguilera-Gomez, A., Vuillez, I., Braakman, I., Hafen, E., Stocker, H. and Rabouille, C. (2015) TORC2 mediates the heat stress response in Drosophila by promoting the formation of stress granules. J Cell Sci, 128, 2497–2508.

59. Riback, J.A., Katanski, C.D., Kear-Scott, J.L., Pilipenko, E.V., Rojek, A.E., Sosnick, T.R. and Drummond, D.A. (2017) Stress-Triggered Phase Separation Is an Adaptive, Evolutionarily Tuned Response. Cell, 168, 1028–1040.

60. Kroschwald, S. and Alberti, S. (2017) Gel or Die: Phase Separation as a Survival Strategy. Cell, 168, 947–948.

61. Berchtold, D., Battich, N. and Pelkmans, L. (2018) A Systems-Level Study Reveals Regulators of Membrane-less Organelles in Human Cells. Molecular Cell, 72, 1035-1049.e1035.

62. Mateju, D., Eichenberger, B., Voigt, F., Eglinger, J., Roth, G. and Chao, J.A. (2020) Single-Molecule Imaging Reveals Translation of mRNAs Localized to Stress Granules. Cell, 183, 1801–1812.

63. Formicola, N., Vijayakumar, J. and Besse, F. (2019) Neuronal ribonucleoprotein granules: Dynamic sensors of localized signals. Traffic, 20, 639–649.

64. Kiebler, M.A. and Bassell, G.J. (2006) Neuronal RNA granules: movers and makers. Neuron, 51, 685–690.

65. Trcek, T. and Lehmann, R. (2019) Germ granules in Drosophila. Traffic, 20, 650–660.

66. Bose, M., Mahamid, J. and Ephrussi, A. (2021) Liquid-to-solid phase transition of RNP granules is essential for their function in the Drosophila germline. bioRxiv, 2021.2003.2031.437848.

67. Aoki, S.T., Lynch, T.R., Crittenden, S.L., Bingman, C.A., Wickens, M. and Kimble, J. (2021) C. elegans germ granules require both assembly and localized regulators for mRNA repression. Nat. Commun., 12, 996.

68. Lee, C.-Y.S., Putnam, A., Lu, T., He, S., Ouyang, J.P.T. and Seydoux, G. (2020) Recruitment of mRNAs to P granules by condensation with intrinsically-disordered proteins. eLife, 9, e52896.

